# Tolerance of liver fluke infection varies between breeds and producers in Scottish beef cattle

**DOI:** 10.1101/2020.07.29.226894

**Authors:** Adam D. Hayward, Philip J. Skuce, Tom N. McNeilly

**Affiliations:** Moredun Research Institute, Pentlands Science Park, Bush Loan, Penicuik, Midlothian, EH26 0PZ, UK

**Keywords:** Liver fluke, *Fasciola* spp., tolerance, disease, productivity

## Abstract

Liver fluke (*Fasciola* spp.) are important helminth parasites of livestock globally and cause significant reductions in health and productivity of beef cattle. Attempts to control fluke have been thwarted by the difficulty of vaccine design, the evolution of flukicide resistance, and the need to control the intermediate snail host. Mechanisms to reduce the impact of parasites on animal performance have typically focused on promoting host resistance – defined as the ability of the host to kill and remove the parasite from its system – and such strategies include improving protein nutrition or selectively breeding for resistance. Organisms, however, have another broad mechanism for mitigating the impact of parasites: they can show tolerance, defined as the ability to maintain health or performance under increasing parasite burden. Tolerance has been studied in the plant literature for over a century, but there are very few empirical studies of parasite tolerance in livestock. In this study, we used data collected from >90,000 beef cattle to estimate the impact of the severity of liver fluke infection on performance and variation in tolerance of fluke. Severity of liver fluke infection was estimated using liver “fibrosis score” on a scale of 0-3 and performance estimated as (1) age at slaughter and (2) daily dead weight gain. Animals with higher fibrosis scores were slaughtered around two weeks later than animals with no fluke, and gained around 10g less weight per day. There was also considerable variation in these effects of fibrosis score, such that animals from different producers and breeds varied in their tolerance of fluke infection. While breeds did not vary in the association between fibrosis and age at slaughter, there was considerable variation among producers: high fibrosis score delayed slaughter by up to 50 days in some producers, but not at all in others. Meanwhile, there was support for variation in the slope of daily dead weight gain on fibrosis score among both breeds and producers, with some unaffected by high fluke scores and some breeds and producers experiencing a 20g/day lower weight gain under high fluke scores. Our results point to the potential for both environmental and genetic variation in tolerance of liver fluke in cattle, paving the way for quantitative genetic and nutritional research into the feasibility of promoting tolerance as a disease mitigation strategy.

**Implications:** Promoting tolerance of disease could help mitigate the impact of disease on livestock productivity, but little research has explored variation in tolerance of livestock diseases or the possibility of promoting tolerance as a mitigation strategy. We used abattoir data to demonstrate that beef cattle vary in their tolerance of fluke infection: while animals from some breeds and some producers experience no impact of fluke on production, others show a large negative effect. Thus, promoting tolerance through management and/or selective breeding could offer a means of reducing the impact of liver fluke on cattle performance.

## Introduction

Liver flukes (*Fasciola* spp.) are among the most important helminth parasites of domestic sheep and cattle worldwide, causing large financial losses (Schweizer, *et al*., 2005) as a result of reduced weight gain (Genicot, *et al*., 1991), milk yield (May, *et al*., 2020), and fertility (May, *et al*., 2019). Control of liver fluke is difficult, with no commercially viable vaccine yet developed (Molina-Hernández, *et al*., 2015), increasing resistance to common flukicides (Kamaludeen, *et al*., 2019), and the necessity of considering the biology of the mud snail (*Galba truncatula*) intermediate host in which clonal amplification of the parasite occurs (Beesley, *et al*., 2018). As such, novel control strategies are likely to be needed in the relatively near future.

Infected hosts have two broad strategies for mitigating the impact of infection upon their health and fitness. One is resistance to infection, defined as the ability of the host to reduce the establishment rate of the parasite and kill and/or remove it from its system (Råberg, *et al*., 2009). Measuring resistance in individual animals is generally straightforward and usually focuses on a measure of infection burden such as helminth faecal egg count (FEC) and other measures of pathogen load, or quantifying pathogen-specific antibody responses. Individuals, genotypes, or breeds with lower pathogen burden or higher antibody levels are generally defined as more resistant. Such measures have been shown to have a considerable heritable component for gastrointestinal nematodes and consequently, breeding for resistance is possible and has been widely implemented with success (Bishop, 2012b). It is clear, however, that potential trade-offs exist between resistance and production traits (Rauw, *et al*., 1998), and enhanced resistance may potentially result in the evolution of counter-measures by the parasite, leading to evasion of host resistance mechanisms, enhanced reproductive rate, or increased parasite virulence (Rausher, 2001).

A second defence mechanism, which has received considerably less attention in the veterinary literature, is tolerance of infection, defined as the ability of the host to maintain health or fitness as parasite burden increases (Råberg, *et al*., 2009). This is quite distinct from resilience to infection, defined as the ability of a host to thrive when infected (Bishop, 2012a), which is actually a product of both resistance and tolerance; indeed many studies purporting to study tolerance are in fact studying resilience (Sakkas, *et al*., 2018). Tolerance has been exceptionally well-studied in the plant literature (Fineblum and Rausher, 1995), with the term being coined to refer to an ability to cope with disease over a century ago (Cobb, 1894). The statistical framework for studying variation in tolerance as “reaction norms” – i.e. variation between groups or genotypes in the rate of change of health or fitness as a function of parasite burden – was also developed in the plant literature (Simms, 2000). Variation in tolerance of vertebrates to pathogens has been more recently demonstrated in both laboratory (Råberg, *et al*., 2007) and wild animal populations (Hayward, *et al*., 2014; Knutie, *et al*., 2017). The benefits of promoting or selecting for tolerance are recognised in the veterinary literature (Bishop, 2012a) and the statistical framework has been described (Doeschl-Wilson, *et al*., 2012), but little empirical work has been undertaken to quantify tolerance variation or explain it in a veterinary setting. A notable exception is tolerance of Porcine Reproductive and Respiratory Syndrome Virus (PRRSV) in pigs, where variation in tolerance has been demonstrated and a candidate tolerance locus identified (Lough, *et al*., 2017; Lough, *et al*., 2018).

Tolerance is most likely to be an important defence mechanism where the parasite is prevalent and resistance is relatively low (Bishop, 2012a), a description which fits the case of liver fluke in cattle. The development of drug resistance in pathogens is inevitable, meaning that other strategies are require for effective control. Promoting tolerance – as opposed to resistance – could be fruitful because tolerance minimizes the evolutionary counter-response from the parasite (Rausher, 2001). The first steps towards designing tolerance-boosting therapies will be quantifying variation in tolerance and identifying its drivers (Vale, *et al*., 2016). Here, we use data collected from slaughtered cattle and use random regression modelling to estimate variation in tolerance as a measure of liver fluke infection between breeds and producers. Our results demonstrate the potential for both genetic and environmental factors to drive variation in tolerance of this important parasite.

## Materials and methods

### Data

The data used in this study were provided by Scotbeef Ltd., Scotland’s largest red meat producers, and were collected between February 2^nd^ 2018 and February 1^st^ 2019. This routinely-collected dataset included information on the identity of the producer, and the breed, sex, date of birth and age at slaughter (in days) of each animal. Carcass data collected included weight, grading, conformation score, fatness; daily dead weight gain was calculated as carcass weight divided by age at slaughter in days. There were also data on whether or not each animal had received a treatment for liver fluke, and the date that the treatment was administered, although information on the product, active compound and dose rate were not available. Livers were inspected for liver fibrosis and assigned a score between 0 (no evidence of fibrosis) and 3 (severe fibrosis), described recently as a proxy for the severity of liver fluke infection (Mazeri, *et al*., 2017). The full dataset consisted of 92,119 animals from 141 breeds and 884 producers.

### Statistical analysis

First, we assessed the overall association between liver fibrosis and two performance traits that were analysed separately: age at slaughter and daily dead weight gain. For each trait, we fitted linear mixed-effects models using the R package ‘glmmTMB’ (Brooks, *et al*., 2017) with breed and producer as random effects and the sex of the animal, whether or not it had been treated for liver fluke, and liver fibrosis score as fixed categorical variables. We compared this model to a model without fibrosis score using a likelihood ratio test (LRT) in order to determine whether fibrosis score was significantly associated with performance. Once missing values were removed, we analysed 91,683 animals from 113 breeds and 875 producers for age at slaughter, and 92,058 animals from 114 breeds and 884 producers for daily dead weight gain.

Next, we assessed whether the change in performance with fibrosis score varied between breeds and/or producers by applying a “reaction norm” approach using random regression models. The approach works on the basis that tolerance is measured as the slope of some measure of performance on disease burden, and testing for variation between groups or individuals in that slope (Simms, 2000; Doeschl-Wilson, *et al*., 2012). For each trait, we first fitted a LMM in ‘glmmTMB’ with breed and producer as random effects, sex and fluke treatment as categorical fixed effects, and fibrosis score as a continuous fixed effect, standardized to be between −1 and +1 (model 1). We then fitted models of the same structure, but with interactions between standardized fibrosis score and the random effects of breed (model 2) or producer (model 3) or both breed and producer (model 4). We tested the significance of the random slope terms by comparing model 2 and 3 with model 1, and by comparing model 4 with models 2 and 3 using LRTs.

## Results

Fibrosis score was significantly associated with age at slaughter, with animals with scores of 1, 2, and 3 taking 13.05±1.82SE, 16.86±2.18 and 14.58±2.15 more days, respectively, to reach slaughter weight than animals with a fibrosis score of 0 (LRT: χ^2^=215.29, DF=3, p<0.001; Figure 1A). Males were slaughtered around a week earlier than females (estimate = −6.79±0.85, χ^2^=63.09, DF=1, p<0.001) and animals that had ever received a fluke treatment took around 3 weeks longer to reach slaughter (estimate = 22.64±1.82, χ^2^=154.04, DF=1, p<0.001). Similarly, a non-zero fibrosis score was associated with lower daily dead weight gain, with animals with fibrosis scores of 1, 2 and 3 gaining −10.0±0.8SE, −12.5±1.6 and −10.1±1.6 fewer grams per day, respectively (LRT: χ^2^=214.81, DF=3, p<0.001; Figure 1B). Males had greater daily dead weight gain than females (estimate = 59.5±0.6g/day, χ^2^=8139.8, DF=1, p<0.001) and animals that had been treated for fluke gained less weight per day (estimate = −14.0±1.4g/day, χ^2^=102.46, DF=1, p<0.001).

**Figure 1.**
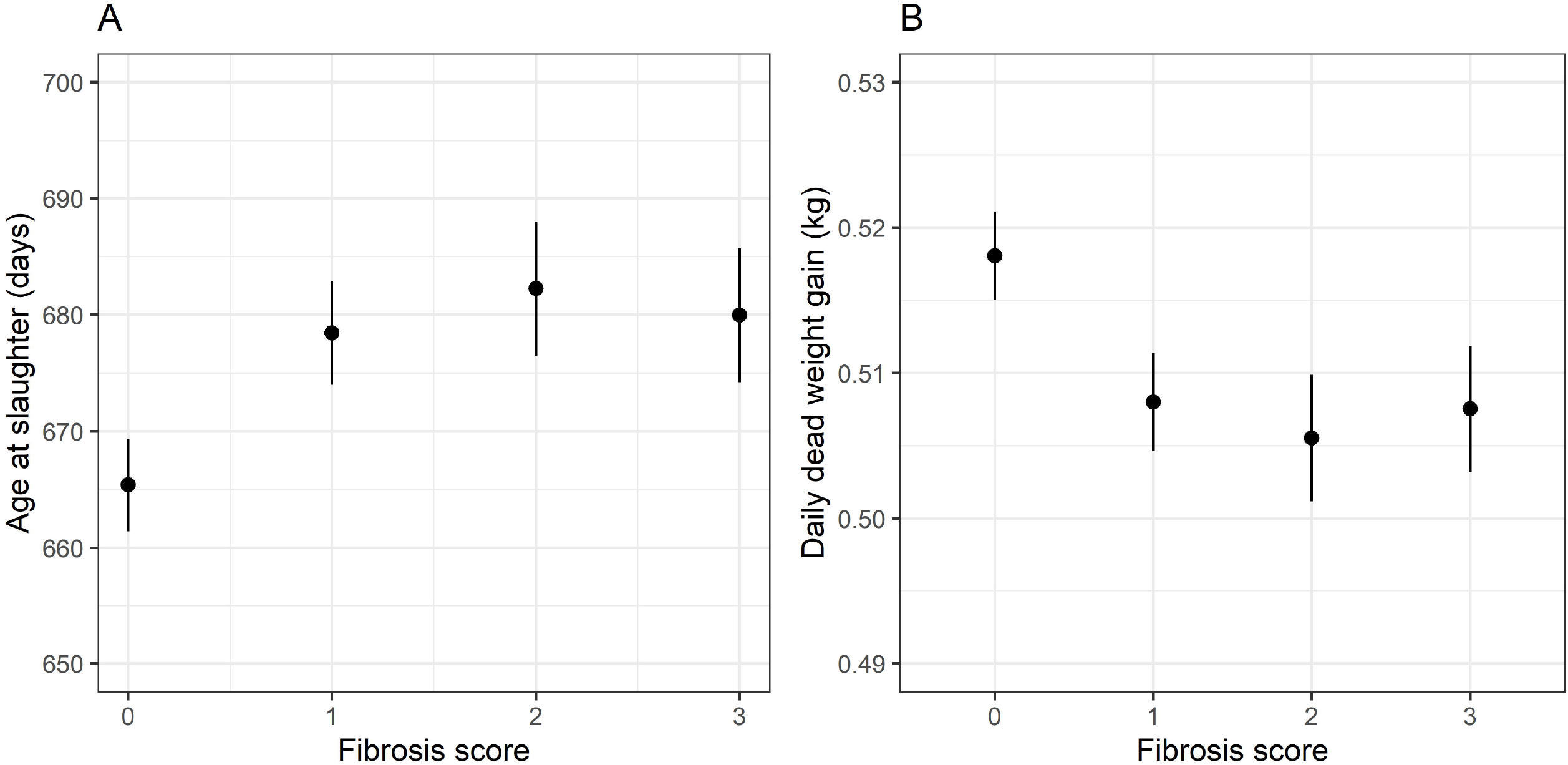
Associations between fibrosis score and (A) age at slaughter and (B) daily dead weight gain, showing means and 95% confidence intervals estimated by linear mixed-effects effects models.

The results of the random regression models for age at slaughter are shown in Table 1. There was no support for a random slope of breed-by-fibrosis (model 2; Figure 2A), but there was evidence to support variation in the slope of age at slaughter on fibrosis between producers (model 3; Figure 2C), which held in the presence of the random slope of breed-by-fibrosis (compare model 4 to model 2). The average estimated delay in age at slaughter between an animal with a fibrosis score of 3 compared to a fibrosis score of 0 was approximately 18 days; while there was little variation around this between breeds (Figure 2B), there was substantially more among producers (Figure 2D), with some producers showing negligible differences in age at slaughter with increasing fibrosis score, and others showing a delay of 50 or even 60 days.

**Table 1.** A comparison of random regression models testing for variation in tolerance of liver fibrosis between breeds and producers, where the phenotype of interest is age at slaughter. Random effects are B = breed, P = producer, with the interaction with F = fibrosis score denoting random slopes of age at slaughter on fibrosis score. “Comparison” shows which model the model in question was tested against using a likelihood ratio test (LRT).

| Model | Random effects | LogLik | Comparison | $\chi^2$ | DF | P |
| --- | --- | --- | --- | --- | --- | --- |
| 1 | B + P | -558233.7 |  |  |  |  |
| 2 | B*F + P | -558232.3 | 1 | 2.95 | 2 | 0.229 |
| 3 | B + P*F | -558200.9 | 1 | 65.76 | 2 | <0.001 |
| 4 | B*F + P*F | -558199.5 | 2 | 65.55 | 2 | <0.001 |
| 4 | B*F + P*F | -558199.5 | 3 | 2.75 | 2 | 0.252 |

**Figure 2.**
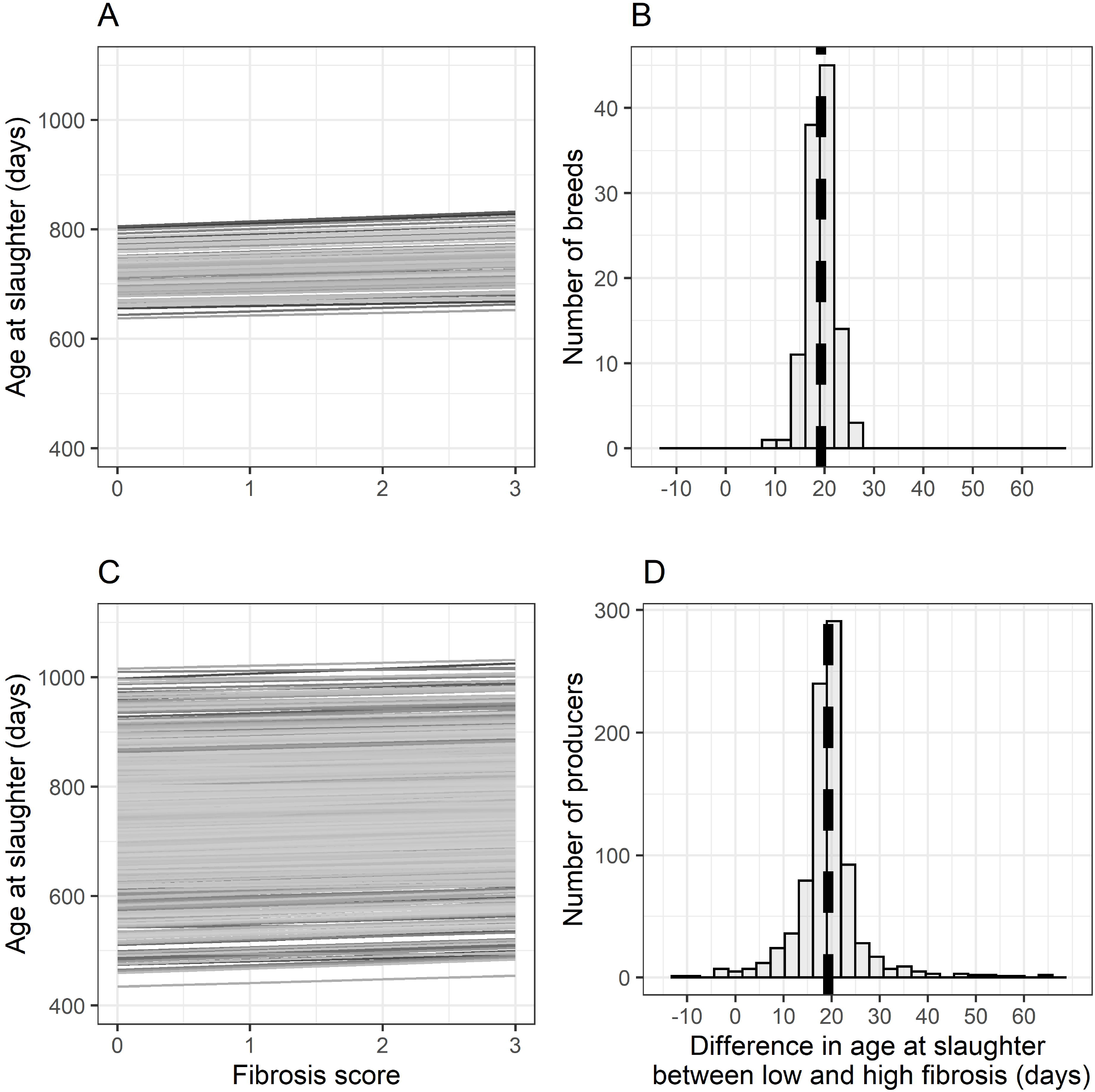
Tolerance variation estimated from random regression models with age at slaughter as the response variable, showing estimated slope of age at slaughter on fibrosis score for each of the (A) breeds and (C) producers, and histograms of the estimated difference in age at slaughter between animals with a fibrosis score of 0 and 3 in different (B) breeds and (D) producers. In B and D, vertical broken line shows model-estimated mean difference in age at slaughter between animals with a fibrosis score of 0 and 3.

The results of the random regression models for daily dead weight gain are shown in Table 2. There was some support for a random slope of breed-by-fibrosis score (model 2; Figure 3A) and stronger support for a random slope of producer-by-fibrosis score (model 3; Figure 3C). However, while the random slope of producer held in the presence of the random slope of breed (compare model 4 with model 2), the converse was not true (compare model 4 with model 3), suggesting that variation in tolerance of fibrosis was more robust among producers than breeds. While the model-estimated average reduction in daily dead weight gain between an animal with a fibrosis score of 3 compared to a fibrosis score of 0 was 0.010kg/day, some breeds and producers showed a difference of zero and hence no effect of fibrosis.

**Table 2.** A comparison of random regression models testing for variation in tolerance of liver fibrosis between breeds and producers, with daily dead weight gain (DDWG) as the performance indicator. Random effects are B = breed, P = producer, with the interaction with F = fibrosis score denoting random slopes of DDWG on fibrosis score. “Comparison” shows which model the model in question was tested against using a likelihood ratio test (LRT).

| Model | Random effects | LogLik | Comparison | $\chi^2$ | DF | P |
| --- | --- | --- | --- | --- | --- | --- |
| 1 | B + P |  |  |  |  |  |
| 2 | B*F + P | 101019.8 | 1 | 7.04 | 2 | 0.030 |
| 3 | B + P*F | 101023.3 | 1 | 66.42 | 2 | <0.001 |
| 4 | B*F + P*F | 101053.0 | 2 | 64.98 | 2 | <0.001 |
| 4 | B*F + P*F | 101055.8 | 3 | 5.60 | 2 | 0.061 |

**Figure 3.**
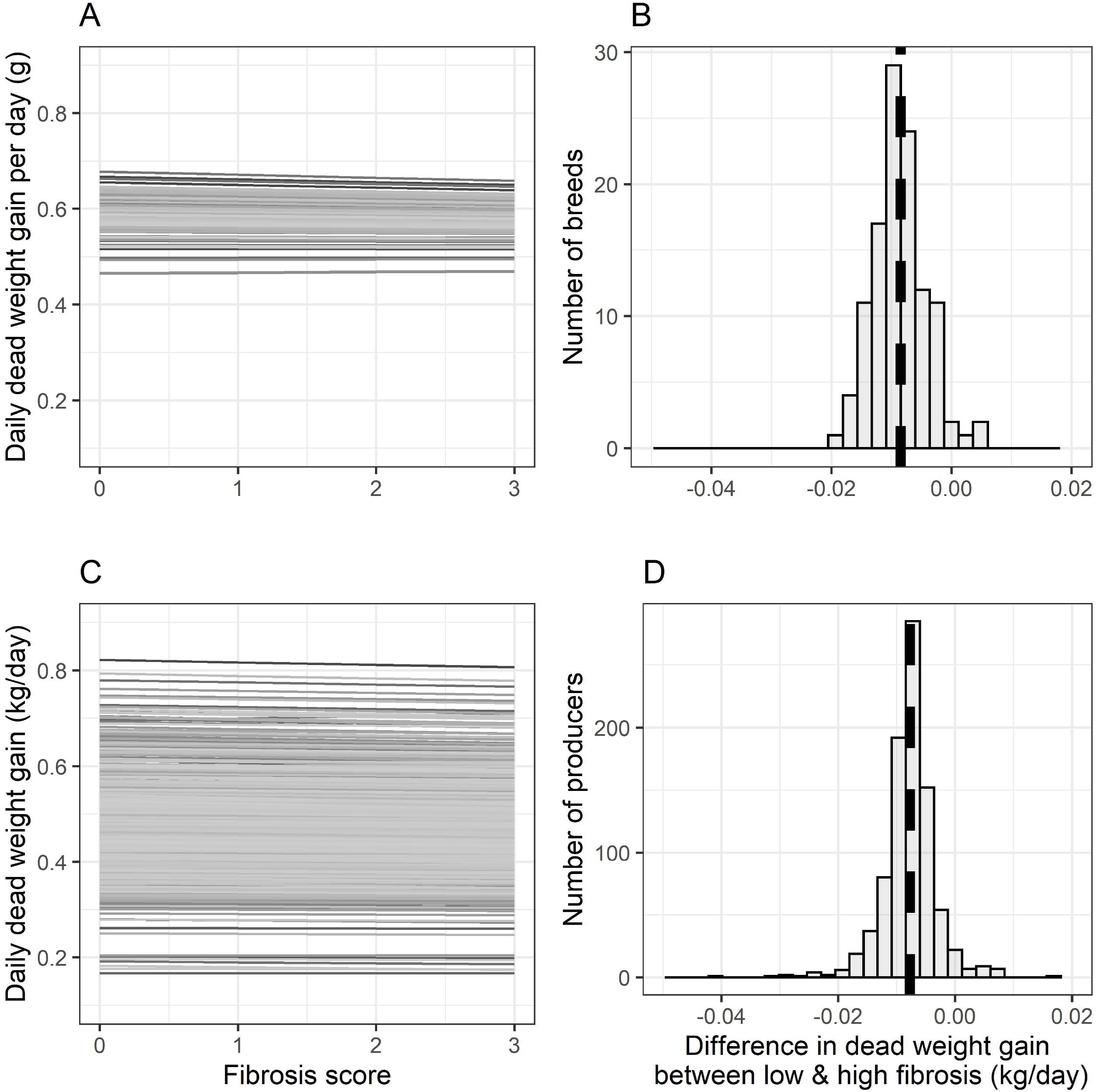
Tolerance variation estimated from random regression models with daily dead weight gain (DDWG) as the response variable, showing estimated slope of DDWG on fibrosis score for each of the (A) breeds and (C) producers, and histograms of the estimated difference in DDWG between animals with a fibrosis score of 0 and 3 in different (B) breeds and (D) producers. In B and D, vertical broken line shows model-estimated mean difference in DDWG between animals with a fibrosis score of 0 and 3.

Meanwhile, some breeds with fibrosis scores of 3 had a difference of up to −0.020kg/day while some animals from some producers with a fibrosis score of 3 had a difference of up to −0.040kg/day.

## Discussion

The results of this study demonstrate the negative impact that liver fluke may have on weight gain in beef cattle resulting in a later age at slaughter. Specifically, we found that cattle with non-zero fibrosis scores gained approximately 10g less per day, and took approximately 2 weeks longer to reach slaughter weight. Previous studies have found similar effects of fluke infection on daily weight gain in beef cattle, although most have used data from experimental infections. These effects range from negligible (Echevarria, *et al*., 1992) to substantial effects of a 0.1kg/day difference between infected and uninfected animals (Jacob, *et al*., 2015) and a difference of 0.7kg/day in Belgian Blue bulls experimentally infected on a feedlot (Genicot, *et al*., 1991). Meanwhile, a previous study using data from the same abattoir as used in the present study – albeit with a smaller sample size of 619 cattle – found substantial effects of fluke infection, with animals with fibrosis scores of 1, 2 and 3 taking on average 34, 93, and 78 days longer to reach slaughter weight, respectively (Mazeri, *et al*., 2017). Most abattoir studies on the impact of fluke in cattle have focused on carcass weight as the performance parameter of interest (Sanchez-Vazquez and Lewis, 2013; Bellet, *et al*., 2016; da Costa, *et al*., 2019), and in these cases differences, while statistically significant, tend to be relatively small, presumably because animals are only sent to slaughter when they reach the requisite target weight.

We then went on to examine whether the linear association between both performance parameters and fibrosis score – our measure of tolerance – varied between breeds and producers. While we did not find variation between breeds in when tolerance was defined in terms of age at slaughter, we did find some support when tolerance was defined by daily dead weight gain. This offers the possibility that genetic variation for tolerance exists, at least at the among-breed level, with some breeds seemingly unaffected by an increasing fibrosis score, and others having considerably lower weight gain. Breeding to mitigate the impact of disease has largely focused on promoting resistance to infection, but the advantages of breeding for tolerance to disease in livestock have also been expounded (Bishop, 2012a; Doeschl-Wilson, *et al*., 2012). These include the fact that tolerance is unlikely to select for pathogens that are better able to evade host resistance (Rausher, 2001; Lough, *et al*., 2017) and that tolerance mechanisms may be general and so offer cross-tolerance to other pathogens (Lough, *et al*., 2017). Further, promoting tolerance is suggested to be potentially advantageous when pathogen prevalence is high, resistance is generally low and elimination has proven difficult due to pathogens evolving in response to treatments (Bishop, 2012a), conditions that apply to liver fluke and gastrointestinal nematodes

We found stronger evidence for variation in tolerance between producers, with stark differences between producers in the effect of fibrosis on both age at slaughter and daily dead weight gain. Such variation could be partly explained by producers rearing different cattle genotypes even within the same breeds, but could be accounted for by a large number of other factors, such as variation in the conditions under which animals are kept (Nakov, *et al*., 2019). Indeed, studies in both wild and lab populations of animals have found variation in tolerance of infection due to variation in diet, including Monarch butterflies (*Danaus plexippus*) feeding on different species of milkweed (Sternberg, *et al*., 2012), BALB/c mice fed on diets of varying protein composition (Clough, *et al*., 2016) and Cuban tree frogs (*Osteopilus septentrionalis*) fed on different resource diets (Knutie, *et al*., 2017). If variation in housing conditions, diet or other management practices are associated with variation in tolerance, it potentially offers a feasible avenue for mitigation of the impacts of fluke infection, although identifying the important factors may be difficult. There is also likely to be variation between fluke genotypes in their life-history traits and virulence (Fairweather, 2011), and so it may be the case that variation in the parasite is largely responsible for the observed variation.

Two further caveats are apparent when considering our results. The first is that, although liver fibrosis score may be a reasonable proxy for fluke burden (Mazeri, *et al*., 2016), it is relatively low resolution, is subjective, and does not distinguish between active and historic infection. Furthermore, liver inspection was shown to be the least effective method out of the five tested at identifying active fluke infection (Mazeri, *et al*., 2016). Nevertheless, the fibrosis score does offer a relatively rapid assessment of fluke infection with some quantitative value on the processing line. The second apparent caveat is that, although the association between fibrosis score and performance in the population as a whole was not linear (**Figure 1**), we imposed a linear association in our random regression models.

In summary, our results show evidence for striking variation both between breeds and producers in their tolerance of a measure of liver fluke burden, offering the possibility of both genetic and environmental variation in tolerance of an important parasite of livestock. Future studies should build on these results in a number of ways. First, studies should aim to use methods – or study parasite species – that enable more reliable estimation of parasite burden in live animals. This will enable, second, repeated measures of parasite burden and performance in the same animal, allowing the estimation of between-individual variation in tolerance. This will facilitate, third, the use of pedigree-based or other methods to estimate genetic variation in tolerance of infection. Once this has been established, potential mechanisms of tolerance variation may be explored. Improved understanding of tolerance could offer new avenues for mitigating the impact of disease on performance (Vale, *et al*., 2016), potentially including genetic improvement programs or adopting management, environmental or nutritional programs that boost animal tolerance.

## Acknowledgements

The authors would like to thank Scotbeef Ltd for providing data, and particularly Suzie England, Uel Morton, William Rowe, Eck Gordon and Sheena Mather. ADH is funded by a Moredun Foundation Research Fellowship. PJS and TNMcN receive funding from the Scottish Government Rural Affairs, Food and the Environment (RAFE) Strategic Research Portfolio 2016-2021.

## Ethics statement

Ethics committee approval was not obtained for this study as the data were obtained from an existing database maintained by Scotbeef Ltd.

## Software and data repository resources

The data are owned by Scotbeef Ltd and as such have not been deposited in an official repository.

## References

Beesley NJ, Caminade C, Charlier J, Flynn RJ, Hodgkinson JE, Martinez-Moreno A, Martinez-Valladares M, Perez J, Rinaldi L and Williams DJL 2018. *Fasciola* and fasciolosis in ruminants in Europe: Identifying research needs. Transboundary and Emerging Diseases 65, 199–216. doi: 10.1111/tbed.12682.

Bellet C, Green MJ, Vickers M, Forbes A, Berry E and Kaler J 2016. *Ostertagia* spp., rumen fluke and liver fluke single- and poly-infections in cattle: An abattoir study of prevalence and production impacts in England and Wales. Preventive Veterinary Medicine 132, 98–106. doi: https://doi.org/10.1016/j.prevetmed.2016.08.010.

Bishop S 2012a. A consideration of resistance and tolerance for ruminant nematode infections. Frontiers in Genetics 3, 168. doi: 10.3389/fgene.2012.00168.

Bishop SC 2012b. Possibilities to breed for resistance to nematode parasite infections in small ruminants in tropical production systems. animal 6, 741–747. doi: doi:10.1017/S1751731111000681.

Brooks ME, Kristensen K, Benthem KJv, Magnusson A, Berg CW, Nielsen A, Skaug HJ, Mächler M and Bolker BM 2017. glmmTMB balances speed and flexibility among packages for zero-inflated Generalized Linear Mixed Modeling. The R Journal 9, 378–400. doi:

Clough D, Prykhodko O and Råberg L 2016. Effects of protein malnutrition on tolerance to helminth infection. Biology Letters 12, 20160189. doi: 10.1098/rsbl.2016.0189.

Cobb NA 1894. Contributions to an economic knowledge of Australian rusts (Uredineae). Agriculture Gazette NSW 5, 239–250. doi:

da Costa RA, Corbellini LG, Castro-Janer E and Riet-Correa F 2019. Evaluation of losses in carcasses of cattle naturally infected with Fasciola hepatica: effects on weight by age range and on carcass quality parameters. International Journal for Parasitology 49, 867–872. doi: https://doi.org/10.1016/j.ijpara.2019.06.005.

Doeschl-Wilson AB, Villanueva B and Kyriazakis I 2012. The first step towards genetic selection for host tolerance to infectious pathogens: obtaining the tolerance phenotype through group estimates. Frontiers in Genetics 3, 265. doi: 10.3389/fgene.2012.00265.

Echevarria FAM, Correa MBC, Wehrle RD and Correa IF 1992. Experiments on anthelmintic control of *Fasciola hepatica* in Brazil. Veterinary Parasitology 43, 211–222. doi: https://doi.org/10.1016/0304-4017(92)90162-3.

Fairweather I 2011. Liver fluke isolates: a question of provenance. Veterinary Parasitology 176, 1–8. doi: https://doi.org/10.1016/j.vetpar.2010.12.011.

Fineblum WL and Rausher MD 1995. Tradeoff between resistance and tolerance to herbivore damage in a morning glory. Nature 377, 517–520. doi: 10.1038/377517a0.

Genicot B, Mouligneau F and Lekeux P 1991. Economic and production consequences of liver fluke disease in double-muscled fattening cattle. Journal of Veterinary Medicine, Series B 38, 203–208. doi: 10.1111/j.1439-0450.1991.tb00862.x.

Hayward AD, Nussey DH, Wilson AJ, Berenos C, Pilkington JG, Watt KA, Pemberton JM and Graham AL 2014. Natural selection on individual variation in tolerance of gastrointestinal nematode infection. PLoS Biology 12, e1001917. doi: 10.1371/journal.pbio.1001917.

Jacob AB, Singh P and Verma AK 2015. Effect of feeding deoiled mahua (*Bassia latifolia*) seed cake on the growth performance, digestibility and balance of nutrients in cross-bred calves during pre-patent period of *Fasciola gigantica* infection. Journal of Animal Physiology and Animal Nutrition 99, 299–307. doi: 10.1111/jpn.12231.

Kamaludeen J, Graham-Brown J, Stephens N, Miller J, Howell A, Beesley NJ, Hodgkinson J, Learmount J and Williams D 2019. Lack of efficacy of triclabendazole against *Fasciola hepatica* is present on sheep farms in three regions of England, and Wales. Veterinary Record 184, 502–502. doi: 10.1136/vr.105209.

Knutie SA, Wilkinson CL, Wu QC, Ortega CN and Rohr JR 2017. Host resistance and tolerance of parasitic gut worms depend on resource availability. Oecologia 183, 1031–1040. doi: 10.1007/s00442-017-3822-7.

Lough G, Rashidi H, Kyriazakis I, Dekkers JCM, Hess A, Hess M, Deeb N, Kause A, Lunney JK, Rowland RRR, Mulder HA and Doeschl-Wilson A 2017. Use of multi-trait and random regression models to identify genetic variation in tolerance to porcine reproductive and respiratory syndrome virus. Genetics Selection Evolution 49, 37. doi: 10.1186/s12711-017-0312-7.

Lough G, Hess A, Hess M, Rashidi H, Matika O, Lunney JK, Rowland RRR, Kyriazakis I, Mulder HA, Dekkers JCM and Doeschl-Wilson A 2018. Harnessing longitudinal information to identify genetic variation in tolerance of pigs to Porcine Reproductive and Respiratory Syndrome virus infection. Genetics Selection Evolution 50, 50. doi: 10.1186/s12711-018-0420-z.

May K, Brügemann K, König S and Strube C 2019. Patent infections with *Fasciola hepatica* and paramphistomes (*Calicophoron daubneyi*) in dairy cows and association of fasciolosis with individual milk production and fertility parameters. Veterinary Parasitology 267, 32–41. doi: https://doi.org/10.1016/j.vetpar.2019.01.012.

May K, Bohlsen E, König S and Strube C 2020. *Fasciola hepatica* seroprevalence in Northern German dairy herds and associations with milk production parameters and milk ketone bodies. Veterinary Parasitology 277, 109016. doi: https://doi.org/10.1016/j.vetpar.2019.109016.

Mazeri S, Sargison N, Kelly RF, Bronsvoort BMd and Handel I 2016. Evaluation of the performance of five diagnostic tests for *Fasciola hepatica* infection in naturally infected cattle using a Bayesian no gold standard approach. PLOS ONE 11, e0161621. doi: 10.1371/journal.pone.0161621.

Mazeri S, Rydevik G, Handel I, Bronsvoort BMd and Sargison N 2017. Estimation of the impact of *Fasciola hepatica* infection on time taken for UK beef cattle to reach slaughter weight. Scientific Reports 7, 7319. doi: 10.1038/s41598-017-07396-1.

Molina-Hernández V, Mulcahy G, Pérez J, Martínez-Moreno Á, Donnelly S, O’Neill SM, Dalton JP and Cwiklinski K 2015. *Fasciola hepatica* vaccine: we may not be there yet but we’re on the right road. Veterinary Parasitology 208, 101–111. doi: 10.1016/j.vetpar.2015.01.004.

Nakov D, Hristov S, Stankovic B, Pol F, Dimitrov I, Ilieski V, Mormede P, Hervé J, Terenina E, Lieubeau B, Papanastasiou DK, Bartzanas T, Norton T, Piette D, Tullo E and van Dixhoorn IDE 2019. Methodologies for assessing disease tolerance in pigs. Frontiers in Veterinary Science 5, doi: 10.3389/fvets.2018.00329.

Råberg L, Sim D and Read AF 2007. Disentangling genetic variation for resistance and tolerance to infectious diseases in animals. Science 318, 812–814. doi:

Råberg L, Graham AL and Read AF 2009. Decomposing health: tolerance and resistance to parasites in animals. Philosophical Transactions of the Royal Society of London B 364, 37–49. doi:

Rausher MD 2001. Co-evolution and plant resistance to natural enemies. Nature 411, 857–864. doi:

Rauw WM, Kanis E, Noordhuizen-Stassen EN and Grommers FJ 1998. Undesirable side effects of selection for high production efficiency in farm animals: a review. Livestock Production Science 56, 15–33. doi: http://dx.doi.org/10.1016/S0301-6226(98)00147-X.

Sakkas P, Oikeh I, Blake DP, Nolan MJ, Bailey RA, Oxley A, Rychlik I, Lietz G and Kyriazakis I 2018. Does selection for growth rate in broilers affect their resistance and tolerance to *Eimeria maxima?* Veterinary Parasitology 258, 88–98. doi: https://doi.org/10.1016/j.vetpar.2018.06.014.

Sanchez-Vazquez MJ and Lewis FI 2013. Investigating the impact of fasciolosis on cattle carcase performance. Veterinary Parasitology 193, 307–311. doi: https://doi.org/10.1016/j.vetpar.2012.11.030.

Schweizer G, Braun U, Deplazes P and Torgerson PR 2005. Estimating the financial losses due to bovine fasciolosis in Switzerland. Veterinary Record 157, 188–193. doi: 10.1136/vr.157.7.188.

Simms E 2000. Defining tolerance as a norm of reaction. Evolutionary Ecology 14, 563–570. doi: 10.1023/a:1010956716539.

Sternberg ED, Lefèvre T, Li J, de Castillejo CLF, Li H, Hunter MD and de Roode JC 2012. Food plant derived disease tolerance and resistance in a natural butterfly-plant-parasite interactions. Evolution 66, 3367–3376. doi: 10.1111/j.1558-5646.2012.01693.x.

Vale PF, McNally L, Doeschl-Wilson A, King KC, Popat R, Domingo-Sananes MR, Allen JE, Soares MP and Kümmerli R 2016. Beyond killing: can we find new ways to manage infection? Evolution, Medicine, and Public Health 2016, 148–157. doi: 10.1093/emph/eow012.

